# BrainStat: a toolbox for brain-wide statistics and multimodal feature associations

**DOI:** 10.1101/2022.01.18.476795

**Authors:** Reinder Vos de Wael, Şeyma Bayrak, Oualid Benkarim, Peer Herholz, Sara Larivière, Raul Rodriguez-Cruces, Casey Paquola, Seok-Jun Hong, Bratislav Misic, Alan C. Evans, Sofie L. Valk, Boris C. Bernhardt

## Abstract

Analysis and interpretation of neuroimaging datasets has become a multidisciplinary endeavor, relying not only on statistical methods, but increasingly on associations with respect to other brain-derived features such as gene expression, histological data, and functional as well as cognitive architectures. Here, we introduce BrainStat - a toolbox for *(i)* univariate and multivariate linear models in volumetric and surface-based brain imaging datasets, and *(ii)* multidomain feature association of results with respect to spatial maps of *post-mortem* gene expression and histology, task-based fMRI meta-analysis, as well as resting-state fMRI motifs across several common surface templates. The combination of statistics and feature associations into a turnkey toolbox streamlines analytical processes and accelerates cross-modal research. The toolbox is implemented in both Python and MATLAB, two widely used programming languages in the neuroimaging and neuroinformatics communities. BrainStat is openly available and complemented by an expandable documentation.

## Introduction

Neuroimaging enables brain-wide measures of morphology, microstructure, function, and connectivity in individuals as well as large cohorts (Casey et al., 2018; Glasser et al., 2013; Royer et al., 2021). Through an increasing array of powerful image processing techniques (Cameron et al., 2013; Esteban et al., 2019; Fischl, 2012; Kim et al., 2005), these data can be brought into a standardized frame of reference including stereotaxic voxel spaces, such as the commonly used MNI152 space (Collins et al., 2003; Dadar et al., 2018), surface-based space such as fsaverage, MNI152-CIVET surfaces, or grayordinates (Fischl, 2012; Glasser et al., 2013; Kim et al., 2005; Lyttelton et al., 2007; Marcus et al., 2011), as well as parcellation schemes (Desikan et al., 2006; Glasser et al., 2016; Gordon et al., 2016; Schaefer et al., 2017). Registering neuroimaging data to a common space allows for the application of statistical analyses, including mass-univariate generalized linear and mixed-effects models that carry out parallel statistical tests at each measurement unit. Usually, such analyses need to be carried out using multiple tools and programs, reducing the reproducibility of workflows, and increasing the risk of human error. With the current paper, we present BrainStat, a unified toolbox to implement these analyses in a cohesive and transparent framework.

The analytical workflows of neuroimaging studies increasingly rely on the availability of previously acquired datasets of other modalities. When mapped to the same reference frame as the neuroimaging measures, these datasets can be used for contextualization of findings and aid in interpretation and validation of results. For example, results may be contextualized within established motifs of the brain’s functional architecture such as intrinsic functional communities based on resting-state fMRI (Yeo et al., 2011) or functional gradients (Margulies et al., 2016), both allowing for the interpretation of findings based on the established brain’s functional architecture. Another common method for contextualization is automated meta-analysis with Neurosynth (Yarkoni et al., 2011), NiMARE (Salo et al., 2020), or BrainMap (Laird et al., 2005). These tools offer the ability to carry out *ad hoc* meta-analyses across potentially thousands of previously published fMRI studies. Correlating a statistical map with a database of brain activation maps related to cognitive terms, so-called meta-analytical decoding, offers a quantitative approach to infer plausible cognitive processes related to a spatial statistical pattern. Finally, *post mortem* datasets of transcriptomics (Hawrylycz et al., 2012) and histology (Amunts et al., 2013) mapped to a common neuroimaging space enable associations of neuroimaging findings with gene expression and histological patterns (Markello et al., 2021; Paquola et al., 2021). Such findings can provide information on molecular and cellular properties in the brain that spatially co-vary with an observed statistical map. By combining these feature association techniques, the functional, histological, and genetic correlates of neuroimaging findings can be inferred. BrainStat provides an integrated decoding engine to run these feature association techniques.

Our toolbox is implemented in both Python and MATLAB, two common programming languages in the neuroimaging research community. BrainStat relies on a simple object-oriented framework to streamline the analysis workflow. The toolbox is openly available at https://github.com/MICA-MNI/BrainStat with documentation available at https://brainstat.readthedocs.io/. We have compartmentalized the toolbox into two main modules: statistics and contextualization (**Figure 1**). In the remainder of this manuscript, we describe how to perform the analyses shown in **Figure 1**.

**Figure 1.**
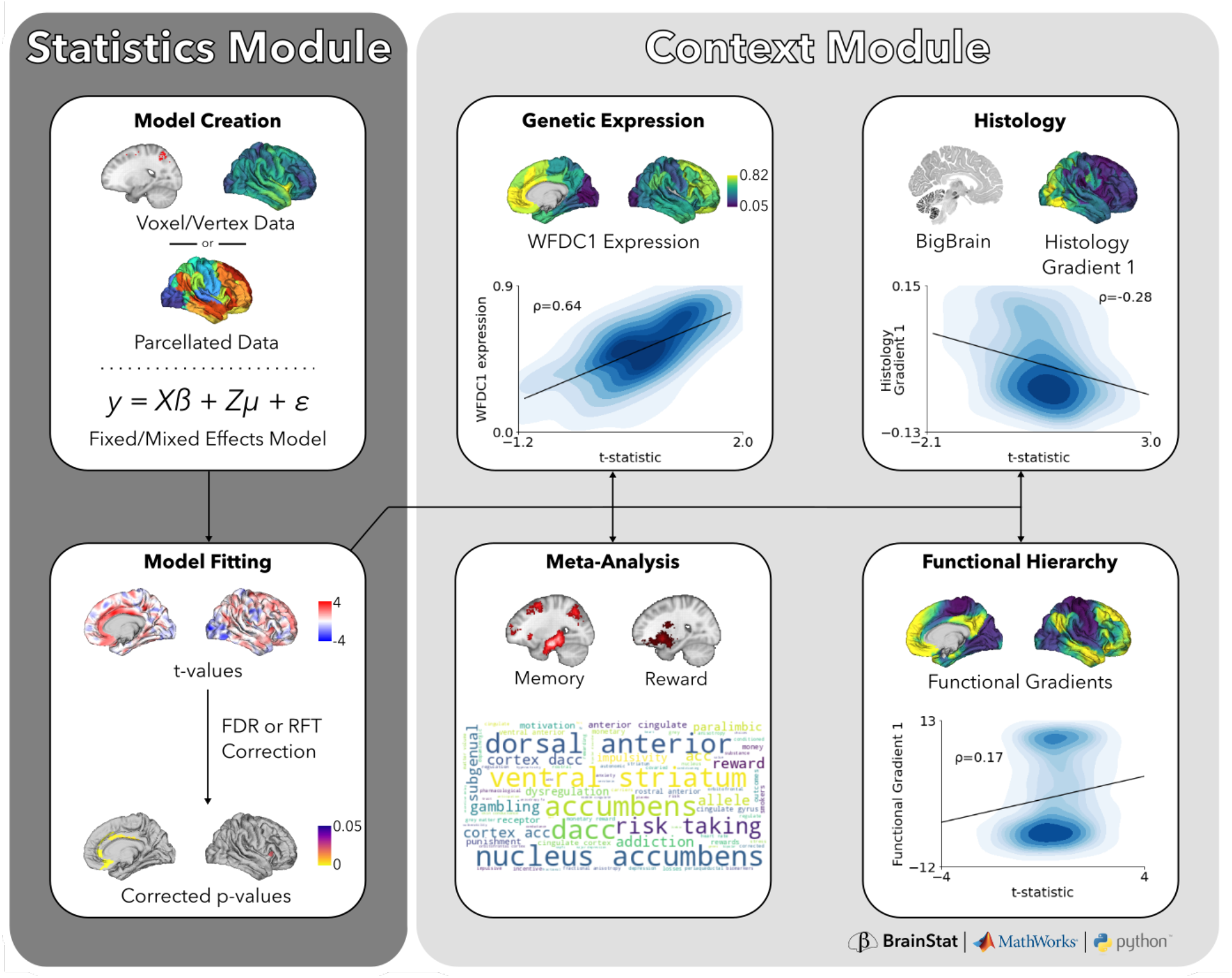
BrainStat workflow. The workflow is split into a statistics module (dark grey) for linear fixed and mixed effects models and a context module (light grey) for contextualizing results with external datasets. To use the statistics module, the user must provide voxel-wise, vertex-wise, or parcel-wise data, and specify a fixed or mixed effects model as well as a contrast probing the model. Once these have been specified, BrainStat computes t-values and corrected p-values for the model. These t-values, or any other brain map, may then be used in the context module to contextualize the statistical results with markers of task fMRI meta-analyses and established functional hierarchies, genetic expression, and histology.

### Statistics Module

The statistics module was built upon a classic, but non-maintained MATLAB package for the flexible implementation of mixed effects models (Worsley et al., 2009). To create and fit such a model, the user provides a subject-by-region-by-variate response matrix as well as a predictor model, created using an intuitive model formula framework. This approach allows for a straightforward definition of fixed/random effects as variables of main interest or control covariates, facilitating both cross-sectional as well as longitudinal analyses. To compare the effects of variables of interest (e.g., healthy/disease, age), a contrast may be specified. BrainStat can handle either univariate or multivariate response data and provides two widely used analytical options for multiple comparison corrections, namely false discovery rate and random field theory. False discovery rate controls the proportion of pointwise (i.e., vertex, voxel, parcel) false positives in the data, whereas random field theory corrects for the probability of reporting a cluster-level false positive.

To illustrate the statistics module, we downloaded cortical thickness and demographic data of 70 participants, 12 of whom were scanned twice, of the MICs dataset (Royer et al., 2021) (**Figure 2A**). We created a linear model with age and sex as well as their interaction effect as fixed effects, and subject as a random effect (**Figure 2B**). These are set up using the FixedEffect and MixedEffect (named as such as it may contain both random and fixed effects) classes. Next, we defined the contrast as -age, i.e., positive t-values denote decreasing cortical thickness with age. This model was fitted on cortical thickness data using a one-tailed test. The t-values, cluster-wise and peak p-values derived from random field theory, as well as the vertexwise p-values derived with a correction for false discovery rate were plotted in **Figure 2C**. We found that, in the MICs dataset, there is evidence for an effect of age on cortical thickness at the cluster level based on random-field theory, but no evidence for significant peaks within these clusters and little significance at a vertex-level. This suggests that the effect covers large regions, rather than local foci. Although we used a liberal cluster-defining threshold (p<0.01) for this illustration, we generally recommend a more stringent threshold (p<0.001), particularly if data with little spatial smoothing are used (Eklund et al., 2016; Woo et al., 2014).

**Figure 2.**
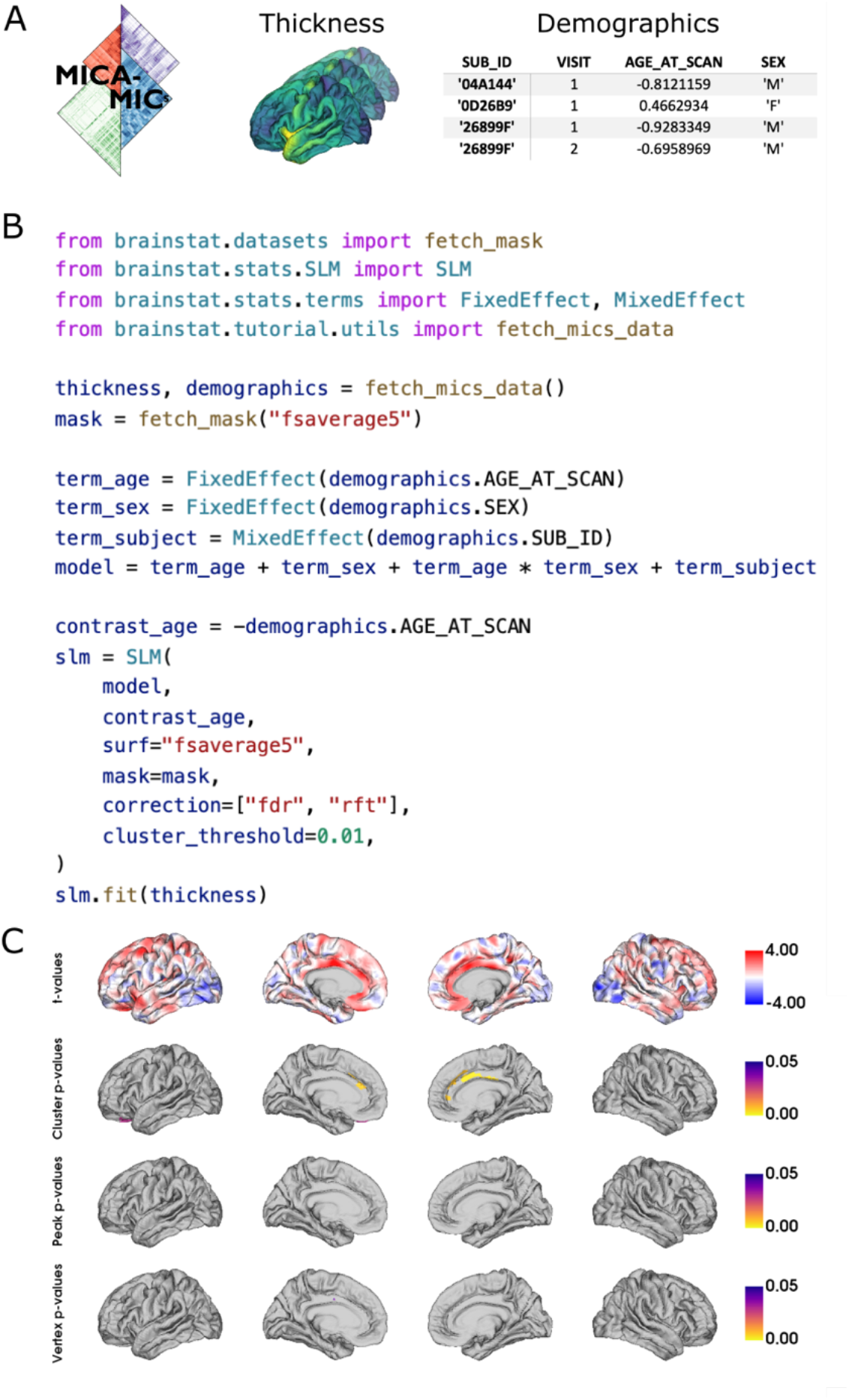
Example Python code for fitting a fixed effect general linear model of the effect of age on cortical thickness with BrainStat. (A) The MICS data included with BrainStat contains cortical thickness and demographics data. The demographics data contains hashed subject IDs (SUB_ID), visit number (VISIT), z-scored age on the day of scanning (AGE_AT_SCAN), and sex (SEX). (B) We create a linear model in the form of *Y* = *intercept* + *age* + *sex* + *age* ∗ *sex* + *random*(*subject*). Note that the intercept is included in the model by default. Third, we initialize the model with an age contrast and request both p-values corrected with random field theory (i.e., “rft”) as well as those corrected for false discovery rate (i.e., “fdr”). Lastly, we fit the model to the cortical thickness data. (C) Negative t-values (blue) indicate decreasing cortical thickness with age, whereas positive t-values (red) depict increasing cortical thickness with age. Significant peak-wise and cluster-wise p-values (p<0.05) are shown for a random field theory (RFT) correction (cluster defining threshold p<0.01) as well as vertex-wise p-values (p<0.05) derived with false discovery rate (FDR) correction. Figure plotting code was omitted for brevity. Python and MATLAB code for this model, as well as code for plotting these figures can be found in the supplemental Jupyter notebook and live script

### Context Module

The context module enables the association of statistical maps with multimodal neural features. As of version 0.3.6, the context module can link to: (*i*) *in-vivo* task-based fMRI meta-analysis, (*ii*) *in-vivo* functional motifs derived from resting-state fMRI, (*iii*) *post-mortem* genetic expression, and (*iv*) *post-mortem* histology/cytoarchitecture (**Figure 1**). The meta-analysis submodule tests for associations between brain maps and task-fMRI meta-analyses associated with particular terms (Salo et al., 2020; Yarkoni et al., 2011). The resting-state module contextualizes neuroimaging findings relative to functional gradients (Margulies et al., 2016; Vos de Wael et al., 2020), a low-dimensional approach to represent the functional connectome. The transcriptomics submodule extracts gene expression from the Allen Human Brain Atlas (Hawrylycz et al., 2012). Lastly, the histological submodule fetches cell-body-staining intensity profiles from the BigBrain (Paquola et al., 2021), a 3D reconstruction of human brain cytoarchitecture (Amunts et al., 2013). These submodules all support common surface templates and, wherever feasible, custom parcellations. Collectively, they pave the way for enrichment analysis of statistical results with respect to aspects of micro- and macroscale brain organization.

#### Meta-analytic decoding

The meta-analytic decoding submodule of BrainStat uses data derived from Neurosynth and NiMARE (Salo et al., 2020; Yarkoni et al., 2011) to decode a statistical map in terms of its cognitive underpinnings. In short, a meta-analytic activation map is created for many (cognitive) terms, and these maps may be correlated to a given statistical map to identify terms with the strongest relationship to the statistical map. This approach allows for the identification of indirect associations to cognitive terms used across a wide-range of previously published task-based functional neuroimaging studies, without relying on cognitive tasks acquired in the same cohort. Indeed, meta-analytic decoding has been used by several groups to evaluate the cognitive associations of their neuroimaging findings [e.g., (Chang et al., 2013; Margulies et al., 2016; Paquola et al., 2019; Vogel et al., 2020; Vos de Wael et al., 2018)].

For each term in the Neurosynth database, we computed which studies used the term with a frequency of at least 1 in 1000 words (default parameter in NiMARE). Next, the meta-analytic maps were computed for these labels using a multilevel kernel density Chi-square analysis implemented in NiMARE (Salo et al., 2020; Wager et al., 2007). For any user-provided surface-based statistical map, we interpolate the map from surface space to volume space. Lastly, for every meta-analytic map in the database, using every voxel that exists within both the meta-analytic map as well as the statistical map we compute a product-moment correlation. An example of retrieving correlations with meta-analytic terms for the t-statistic map derived earlier is shown in **Figure 3**.

**Figure 3.**
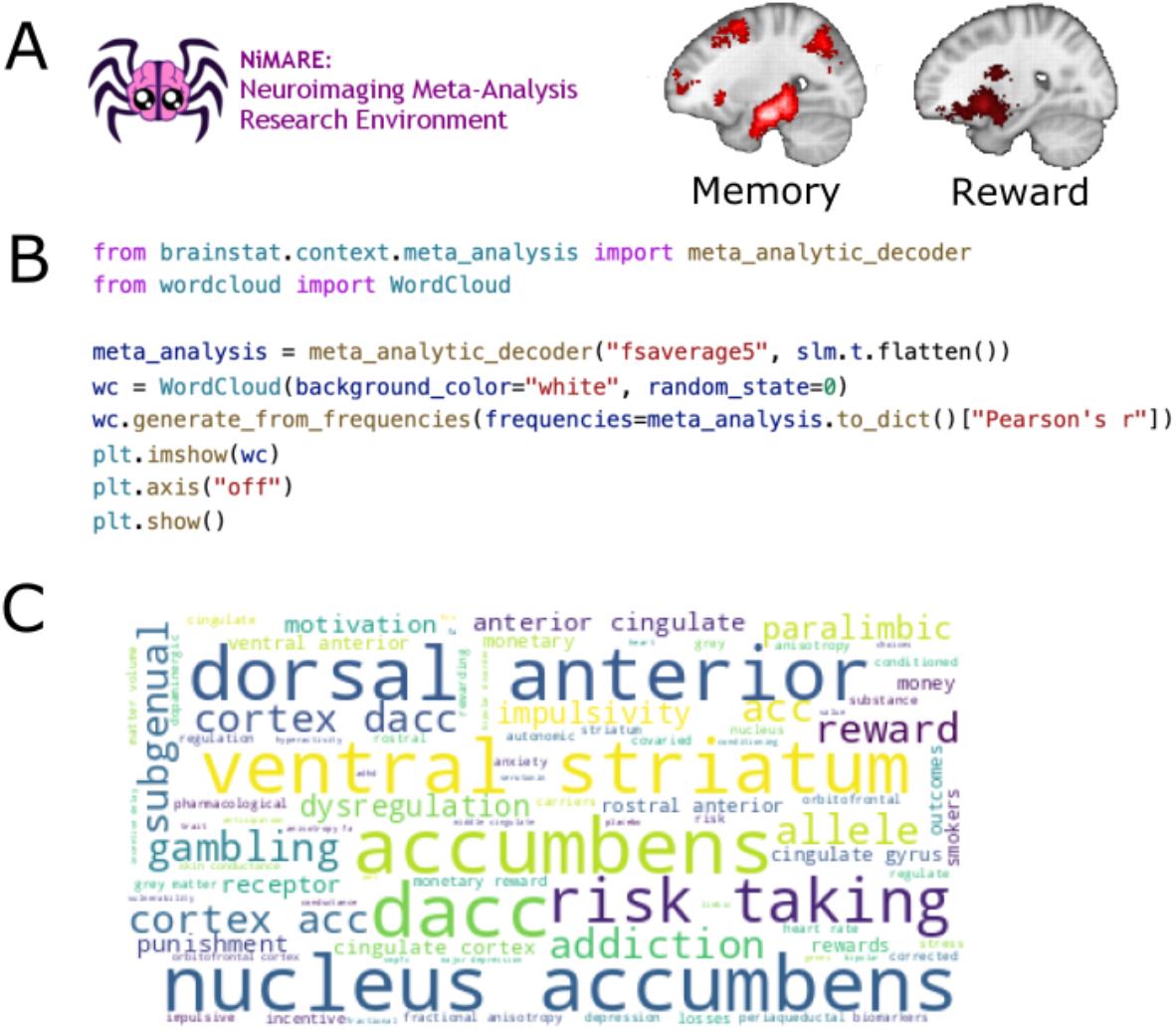
(A) Using the NiMARE toolbox (Salo et al., 2020) and the Neurosynth database (Yarkoni et al., 2011) we derived feature maps for every feature name in the Neurosynth database. We show an example association map for the terms “memory” and “reward”. (B) Example Python code for computing the correlations between the t-statistic map and every term in the Neurosynth database, and plotting these correlations as a word cloud. This code as well as the MATLAB equivalent can be found in the supplemental Jupyter notebook and live script. (C) The resultant word cloud from the code in **Figure 3B**.

#### Resting-state motifs

The functional architecture of the brain at rest has been described both as a set of continuous dimensions, also called gradients; (Margulies et al., 2016; Vos de Wael et al., 2020). These gradients highlight gradual transitions between regions and can be used to embed findings into the functional architecture of the human brain by assessing point-wise relationships with other markers. Indeed, prior studies have used functional gradients to assess the relationship of the brain’s functional architecture to high level cognition (Murphy et al., 2019; Shine et al., 2019), hippocampal subfield connectivity (Vos de Wael et al., 2018), amyloid beta expression and aging (Lowe et al., 2019), microstructural organization (Huntenburg et al., 2017; Paquola et al., 2019), phylogenetic changes (Xu et al., 2020), and alterations in disease states (Caciagli et al., 2021; Hong et al., 2019; Tian et al., 2019).

The functional gradients included with BrainStat were computed by resampling the mean functional connectivity matrix of the S1200 release to fsaverage5 (to reduce computational complexity), and subsequently computing gradients with BrainSpace using the following parameters: cosine similarity kernel, diffusion map embedding, *alpha=0*.*5, sparsity=0*.*9*, and *diffusion_time=0* (Vos de Wael et al., 2020). Example code for computing the correlations between the first functional gradient and the t-statistic map are shown in **Figure 4**. We find a low Spearman correlation (ρ=0.17) between these two maps. However, to test for significance of this correlation we need to correct for the spatial autocorrelation in the data (Markello & Misic, 2021). Three such corrections, namely spin test (Alexander-Bloch et al., 2018), Moran spectral randomization (Wagner & Dray, 2015), and variogram matching (Burt et al., 2020), are included in BrainSpace (Vos de Wael et al., 2020), a dependency of BrainStat.

**Figure 4.**
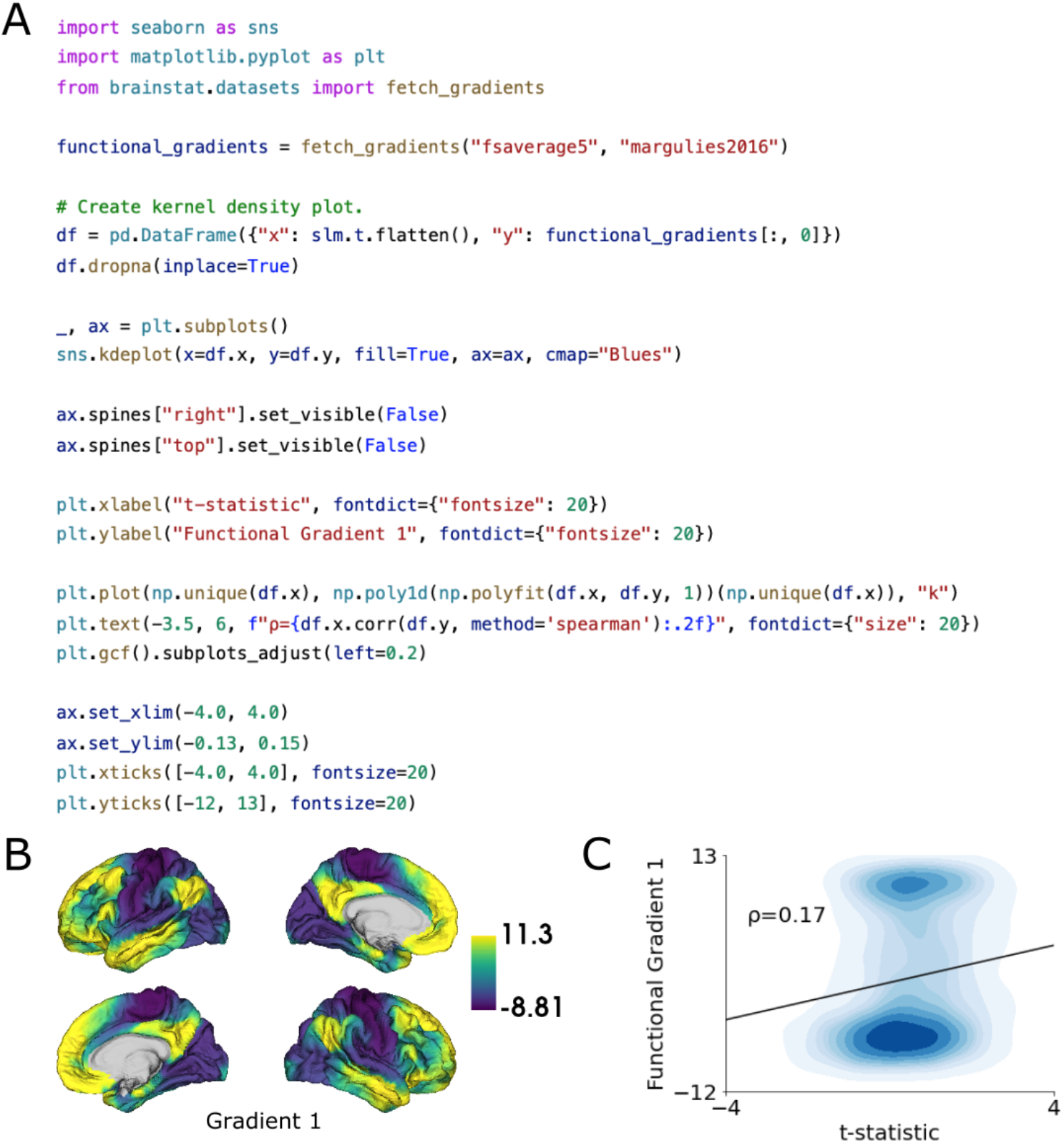
(A) Example Python code for computing and plotting the correlation of the t-statistic map and the first functional gradient. Cortical surface plotting code was omitted for brevity. See the supplemental Jupyter notebook (Python) and live script (MATLAB) for executable versions of this code as well as additional figure building code. (B) First functional gradient plotted to the surface. (C) Kernel density plot of the t-statistic map and the first functional gradient.

#### Genetic Expression

The Allen Human Brain Atlas (Hawrylycz et al., 2012) is a database of microarray gene expression data of over 20,000 genes derived from *post-mortem* tissue samples obtained from six adult donors. This resource may be used to derive associations between neuroimaging data and molecular factors (Arnatkeviciute, Fulcher, Bellgrove, et al., 2021) and thus yield insights into the mechanisms giving rise to anatomical and connectomic markers. For example, such data may be used to study associations between genetic factors and functional connectivity (Cioli et al., 2014; Krienen et al., 2016; Richiardi et al., 2015), anatomical connectivity (Goel et al., 2014; Park, Bethlehem, et al., 2021), as well as alterations of connectivity in disease (Park, Hong, et al., 2021; Romme et al., 2017). The genetic decoding module of BrainStat uses the abagen toolbox (Markello et al., 2021) to compute genetic expression data for a given parcellation. Using default parameters, abagen performs the following procedure. First it fetches and updates the MNI152 coordinates of tissue samples of all six donors using coordinates provided by the *alleninf* package (https://github.com/chrisgorgo/alleninf). Next, it performs an intensity-based filtering of the probes to remove those that do not exceed background noise. Subsequently, for probes indexing the same gene, it selects the probe with the highest differential stability across donors. The tissue samples are then matched to regions in the parcellation scheme. Expression values for each sample across genes and for each gene across samples are normalized using a scaled robust sigmoid normalization function. Lastly, samples within each region are averaged within each donor, then averaged across donors. For details on the procedures with non-default parameters please consult the abagen documentation (https://abagen.readthedocs.io/). In Python, BrainStat calls abagen directly, and as such all parameters may be modified. In MATLAB, where abagen is not available, we included genetic expression matrices precomputed with abagen with default parameters for many common parcellation schemes. In **Figure 5** we show an example of fetching the genetic expression for a previously defined functional atlas (Schaefer et al., 2017), and correlating this to the t-statistic map. The expression derived from this module can then be used in further analyses, for example by deriving the principal component of genetic expression and comparing it to previously derived statistical maps.

**Figure 5.**
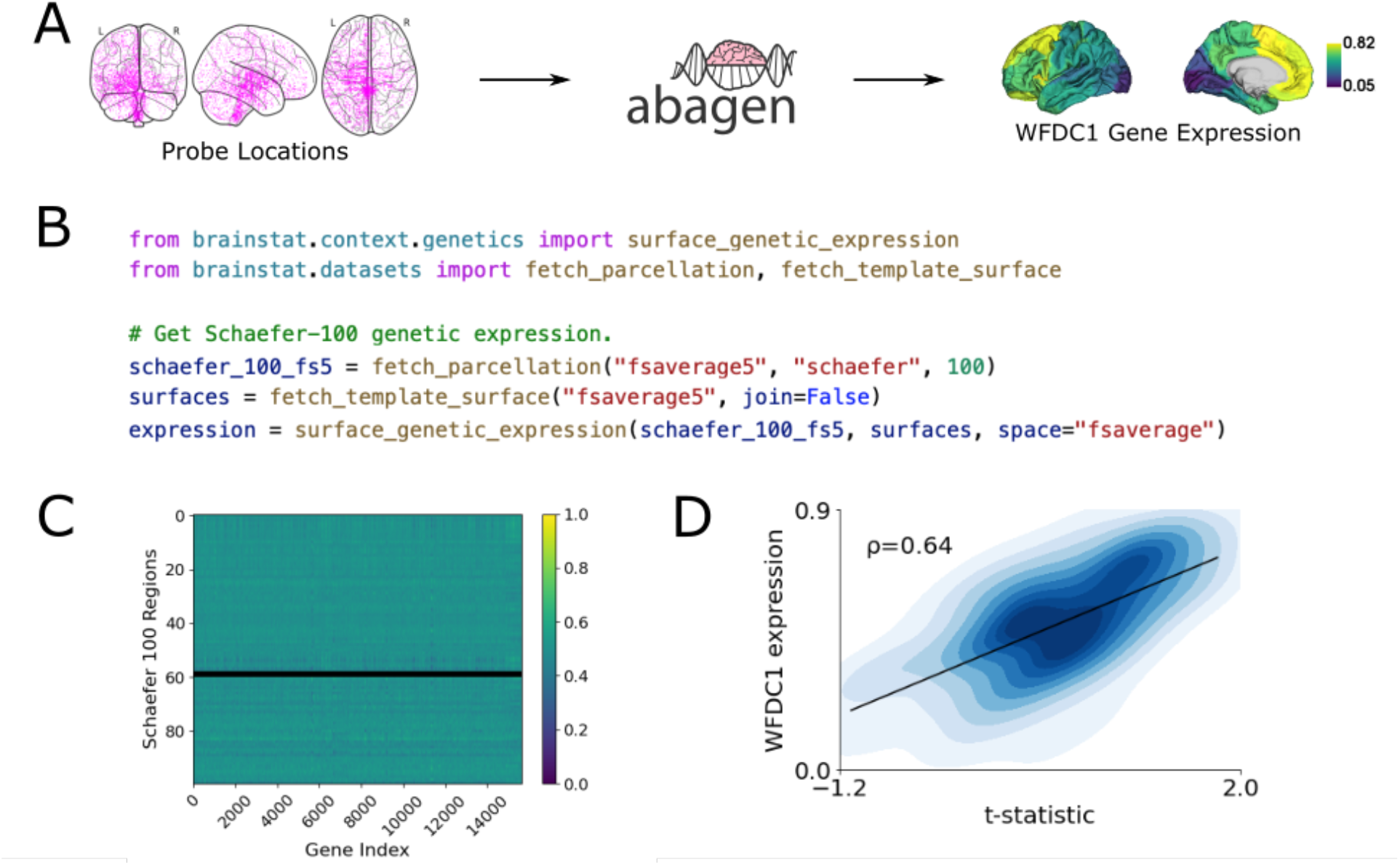
(A) BrainStat uses data from the Allen Human Brain atlas (Hawrylycz et al., 2012), provided by the Allen institute, and processed with abagen (Markello et al., 2021) to derive transcription levels of several thousands of genes. Shown are the locations of all probes, as well as the expression of a single gene (WFDC1) within 100 functionally defined regions (Schaefer et al., 2017). (B) Example Python code for computing the genetic expression based on a surface parcellation. (C) The genetic expression matrix depicts the genetic expression, normalized to a range of [0, 1], of all parsed genes across all parcels. Black rows denote regions without samples. (D) Correlation between the t-statistic map and the WFDC1 gene expression. Figure plotting code was omitted for brevity. See the supplemental Jupyter notebook (Python) and live script (MATLAB) for executable versions of this code as well as additional figure building code.

#### Histology

The BigBrain atlas (Amunts et al., 2013) is a three-dimensional reconstruction of a sliced and cell-body-stained human brain. With a digitised isotropic resolution of 20 micrometers, it is the first openly available whole-brain 3D histological dataset. As such, it is well suited for relating neuroimaging markers to histological properties. This may be used, for example, for cross-validating MRI-derived microstructural findings (Paquola et al., 2019; Royer et al., 2020), defining regions of interest based on histological properties (Sitek et al., 2019), or relating connectomic markers to microstructure (Arnatkeviciute, Fulcher, Oldham, et al., 2021). The histology submodule aims to simplify the integration of neuroimaging findings with the BigBrain dataset. This submodule uses surfaces sampled from the BigBrain atlas at 50 different depths across the cortical mantle (Paquola et al., 2021). Covariance of these profiles, also known as microstructural profile covariance (Paquola et al., 2019), is computed with a partial correlation correcting for the mean intensity profile. Principal axes of cytoarchitectural variation are computed from microstructural profile covariance using BrainSpace with default parameters (Vos de Wael et al., 2020). An example of this is shown in **Figure 6**. We find a correlation between the first eigenvector and the t-statistic map of ρ=-0.28.

**Figure 6.**
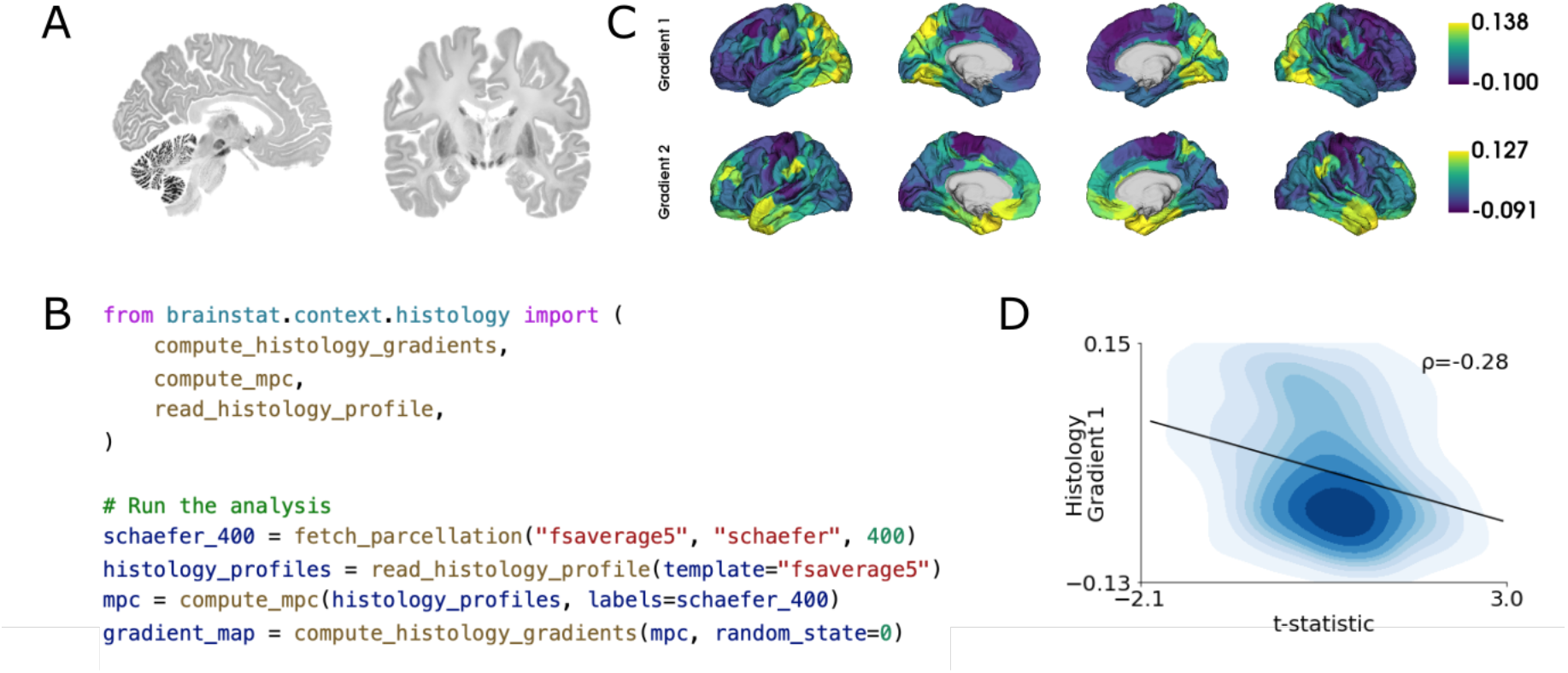
(A) Selected sagittal and coronal slices from the BigBrain atlas. (B) Example Python code for computing gradients of microstructural profile covariance for the Schaefer-400. Histology profiles, as computed in the BigBrainWarp toolbox, are loaded. From these profiles, a microstructural profile covariance matrix is built, and used to compute gradients of histology. (C) Gradients derived from the microstructural profile covariance of the BigBrain atlas. (D) The scatter plot of the t-statistic map and the first gradient of microstructural profile covariance. Figure plotting code was omitted for brevity. See the supplemental Jupyter notebook (Python) and live script (MATLAB) for executable versions of this code as well as additional figure building code.

## Discussion

The analysis of brain imaging data demands tools for uni- and multivariate statistical inference, and increasingly leverages data from multiple resources to facilitate interpretation and contextualization of significant results. Although many tools have been made available by the neuroimaging community to perform individual analytical steps (Larivière et al., 2021; Markello et al., 2021, 2022; Paquola et al., 2021; Salo et al., 2020; Yarkoni et al., 2011), there is currently no package that unifies statistical inference and contextualization approaches. To fill this gap, we presented BrainStat, an integrated toolbox for the univariate and multivariate statistical analysis of brain imaging data and for performing multidomain feature associations to interpret findings. As a fully open access tool implemented in both Python and MATLAB, BrainStat will be easily accessible by the broader neuroscience community. Furthermore, BrainStat is complemented with a thorough and easy-to-follow online documentation, providing even novice users an entry into integrated analyses of brain imaging data.

Linear models are a core technique for neuroimaging-based inference. Indeed, many common univariate and multivariate statistical analyses, including t-tests, F-tests, multiple linear regression, and (M)AN(C)OVA, can be considered special cases of the general linear model (Friston, 2005; Nichols & Holmes, 2002). As such, usage of general linear models is widespread throughout the neuroimaging literature [e.g., (Bernhardt et al., 2018; Clarkson et al., 2011; Truong et al., 2013)]. Despite their prevalence, their implementation is not trivial. BrainStat’s statistics module aims to provide a flexible multivariate linear modeling framework for neuroimaging data (Worsley et al., 2009). The focus of this manuscript was to present the possibilities provided by BrainStat and outline an accessible tutorial for their usage within the toolbox. As such, we demonstrated a mixed effects models for testing the effects of age on cortical thickness. However, given the flexibility of linear models, a large variety of models may be specified within the same framework. Furthermore, the flexible specification of a contrast simplifies testing of fitted models.

Recent years have seen an uptick in the usage of external datasets for the contextualization of MRI-derived results. These datasets may be leveraged for their unique advantages such as the unprecedented spatial resolution of the BigBrain histological atlas (Amunts et al., 2013), the vast number of task-fMRI studies included in the Neurosynth meta-analytical database (Yarkoni et al., 2011), and 3D maps of post-mortem human brain gene expression information aggregated in the Allen Human Brain Atlas (Hawrylycz et al., 2012). These external datasets allow for more comprehensive studies of brain organization and may advance our understanding of fundamental principles of brain organization (Hansen et al., 2021; Larivière et al., 2019). Indeed, prior studies have used these datasets to relate task meta-analyses, genetic expression, and histology to markers of morphology (Valk et al., 2020; Wagstyl et al., 2020; Whitaker et al., 2016), function (Benkarim et al., 2021; Krienen et al., 2016; Margulies et al., 2016; Paquola et al., 2019), and structural connectivity (Romme et al., 2017; Vos de Wael et al., 2021). Though numerous packages exist to enable these analyses (Markello et al., 2021; Paquola et al., 2021; Salo et al., 2020; Yarkoni et al., 2011), these are generally distributed independently and their integration requires expertise, and often proficiency in specific programming languages as there are generally no cross-language implementations. BrainStat brings these tools together into a unified multi-language framework, thereby increasing their accessibility and streamlining the analytics processes of neuroimaging studies. Ultimately, we hope to reduce the barrier to entry of these techniques, reduce the chances of human error, and thereby accelerate cross-modal research in the neuroimaging community.

Theoretical and empirical studies have shown the importance of replicability in science (Ioannidis, 2005; Moonesinghe et al., 2007; Open Science Collaboration, 2015). The proliferation of open-access datasets (Di Martino et al., 2014; Miller et al., 2016; Royer et al., 2021; Van Essen et al., 2013) and software (Fischl, 2012; Marcus et al., 2011; Paquola et al., 2021; Tournier et al., 2012; Vos de Wael et al., 2020) may increase reproducibility by allowing others to redo experiments with identical data and procedures, as well as reducing human error in the analysis (Milham et al., 2018; Poldrack et al., 2017). BrainStat may contribute to this process. By unifying statistical processing and multidomain feature association into a single package across two programming languages, the resulting code will require less customization and technical expertise from the end-user. Furthermore, BrainStat may increase the accessibility to all these methods for researchers in places that lack the institutional expertise to set up such integrated pipelines.

## Code Availability

BrainStat is freely available through PyPi (https://pypi.org/project/brainstat/; installable with *pip install brainstat*), FileExchange (https://www.mathworks.com/matlabcentral/fileexchange/89827-brainstat; installable with the add-on manager in MATLAB), and GitHub (https://github.com/MICA-MNI/BrainStat). Documentation is available at https://brainstat.readthedocs.io/. BrainStat 0.3.6 supports Python 3.7-3.9 and MATLAB R2019b+ on Windows, macOS, and Linux. However, we advise users to consult the installation guide in our documentation for up-to-date requirements.

## Methods

### Tutorial dataset

We studied 70 healthy participants [30 females, age = 31.9 ± 8.9 (mean±SD); 12 of them (5 female) came in for a second visit with age = 32.8 ± 7.5] of the MICS dataset (Royer et al., 2021). Note that these data include subjects not part of the current MICS release. For each visit, two T1w images were derived with the following parameters: MP-RAGE; 0.8mm isotropic voxels, matrix = 320 × 320, 224 sagittal slices, TR = 2300ms, TE = 3.14ms, TI = 900ms, flip angle = 9°, iPAT = 2, partial Fourier = 6/8. Scans were visually inspected for motion artefacts. Processing was performed with micapipe (https://github.com/MICA-MNI/micapipe). In short, cortical surface segmentations were generated from the T1w scans using Freesurfer 6.0 (Fischl, 2012). Subject’s thickness estimates were transformed to the fsaverage5 template and smoothed with a 10mm full-width-at-half-maximum Gaussian kernel.

## Acknowledgments

The authors would like to thank the late Keith Worsley for his invaluable work on the SurfStat toolbox which inspired BrainStat.

## Author Contributions

R.V., and Ş.B. were the lead developers of the toolbox. O.B., P.H., S.V, and B.B. further assisted in toolbox design. R.V., and B.B. drafted the manuscript. All authors assisted in revising the manuscript.

